# Aligned recordings of neural spiking activity and licking behavior in thirsty mice

**DOI:** 10.64898/2026.04.21.720009

**Authors:** Zihang Xu, Binjie Hong, Li Li, Taorong Xie, Zhaorong Chen, Haishan Yao, Tielin Zhang

## Abstract

Electrophysiological data, which serve as a biological signal that bridges neural activity and behavioral tasks, provide an innovative approach to neuroscience research. In this study, we constructed a dataset that contains over 2000 neurons across 117 days recorded in 20 mice containing 28,573 trials. Data for 5 mice were collected from the Secondary Motor Cortex (M2) region 8 mice was derived from the Ventrolateral Striatum (VLS) and 7 mice were from Substantia Nigra pars Reticulata (SNR). We induced licking behavior in head-fixed mice by periodically delivering water through a spout while simultaneously recording spiking activity from three brain regions and behavior related electrical signals. This dataset ensures precise temporal alignment between neural activity and behavioral events, offering a robust foundation for investigating neural encoding mechanisms and simulation of neural activities. This dataset establishes a precise spike-to-event mapping, which enables high decoding accuracy using Multilayer Perceptron (MLP) and Support Vector Machine (SVM). It can serve as a high-quality benchmark for developing encoding and decoding algorithms in neural networks, particularly Spiking Neural Networks (SNNs).

## Background and Summary

Mice have gained increasing prominence in neuroscience and physiology research due to their well-defined genetics [1], controllable behaviors [2] and mature experimental toolkits [3]. For example, the team from the Chinese University of Hong Kong [4] found that contrast gain control may represent a canonical cortical computation and lays a foundation for investigations into the underlying mammal’s auditory mechanisms. In addition, mice were used for the study of classifying electrophysiological and morphological neuron types in the visual cortex [5]. Furthermore, research indicates that the thalamus selectively activates the M1 neurons responsible for the learned movements in skilled individuals [6], underscoring a critical link between the thalamus, M1, and motor learning.

Neural spikes offer significant advantages in studying the mechanisms between neural activity and behavior [7]. As discrete and event-driven signals, spikes enable the temporal precision of spike transmission [8]. This sparse coding strategy enhances computational efficiency in artificial neural systems using deep learning algorithms [9]. For example, Simplicial Convolutional Recurrent Neural Network (SCRNN) has been proposed as a method for decoding the direction of the head from the spikes of navigation cells [10]. Researchers also demonstrated that recording only a small part of the total neural spike responses from the dorsal premotor cortex can help decode 3D hand movement trajectories with high precision [11].

In our dataset, both the electrophysiology data and the licking behavior data are converted to spiking-based data (only valuing 0 or 1), which consists of discrete timestamps marking the precise moments when neurons generate action potentials. We provide data of 20 individual mice that cross 119 days from 3 brain regions, including Secondary Motor Cortex (M2), Substantia Nigra pars Reticulata(SNR) and Ventral Lateral Striatum(VLS). Researchers may utilize the data to investigate the relationship between behavior and neural response on days and subjects. In addition, data collected from different regions facilitate comparative biological research on functional differences between brain regions, including M2, SNR, and VLS.

## Methods

### Surgery

Mice were anesthetized using an intraperitoneal cocktail of fentanyl (0.05 mg/kg), medetomidine (0.5 mg/kg), and midazolam (5 mg/kg), then secured in a stereotaxic frame. Following local lidocaine application, bilateral craniotomies (∼ 1 mm diameter) were performed in the target cortical regions. Viral vectors (2–3 × 10^12^ particles/mL) were delivered via a glass pipet (15– 20 *µ*m tip) using a syringe pump. Nine mice were divided into three experimental groups targeted anterior lateral motor cortex(ALM) [12] inhibition: VGAT-ChR2 [13] mice (*n* = 3), VGAT-Cre mice with AAV-FLEX-ChrimsonR (*n* = 3), and C57BL/6 mice [14] with AAV-CaMKII*α*-GtACR2 (*n* = 3). Control C57BL/6 mice received AAV-hSyn-eGFP. Additional groups received inhibitory opsins in central M2 (*n* = 6, AAV-eNpHR3.0) or medial prefrontal cortex(mPFC) (*n* = 5, AAV-Jaws). Each injection (200–400 nL) was delivered at specific coordinates and depths. Post-injection, optical fibers (200 *µ*m, NA 0.37) were implanted at appropriate positions. We injected 400–500 nL of AAV into VLS at a depth of 3.2 mm and into dorsal striatum(DS) at a depth of 2.0 mm. DS serves as a key integration hub for cortical and thalamic inputs and provides efferent projections to the SNR region [15]. Consequently, targeted modulation of the DS allows for precise functional interrogation of neural dynamics in the SNR. A headplate was affixed with dental cement and mice recovered for 10 days with Rimadyl analgesia before behavioral training.

### Behavioral task

Mice underwent a 48-hour water restriction period before behavioral training. During experiments, the animals were head fixed in acrylic restraint tubes and trained to lick a 3 mm anterior and 1 mm ventral spout. Licking behavior was detected using custom electrical contact sensors [16] or infrared beam-break systems (the latter used during electrophysiological recordings). Following initial habituation (1–2 days of handling and syringe water licking training with 300–500 nL rewards) and free-drinking adaptation (1–2 days of head-fixed sessions with 2–3 *µ*L water delivered every 4 seconds), mice were trained on a fixed-interval licking task (10–30 days) where they received 2–3 *µ*L of 10% sucrose every 10 seconds. Upon demonstrating consistent anticipatory licking, probe trials (3–4× longer than reward intervals, constituting ∼10% of trials and always preceded by 4–14 reinforced trials, which is used to determine whether they have truly mastered the temporal interval pattern) were introduced. Training continued for 20–50 days until probe trial licking patterns (assessed via PSTH) matched reinforced trials, with daily 1-hour sessions in the final phase. For electrophysiology experiments, mice underwent an additional 14– 21 days of training, including 1-hour pre-task head fixation periods to accommodate subsequent cortical electrode implantation. After being trained in the final task phase for 2–3 days, the mice were used for electrophysiological experiments.

The whole experiment procedure can be shown as Fig.1. The experimental protocol is divided into three distinct segments: an interval allowing the mice to rest, a subsequent period for the water-licking task, and a final phase for system reset and initialization. The combined duration of the interval time and the fixation time is approximately 7 seconds. During the experiment, the mice spend about 3 seconds licking water, making the total time for one trial approximately 10 seconds. During the experiment, the behavioral signals of the mice and the corresponding electrophysiological signals are collected synchronously.

**Figure 1:**
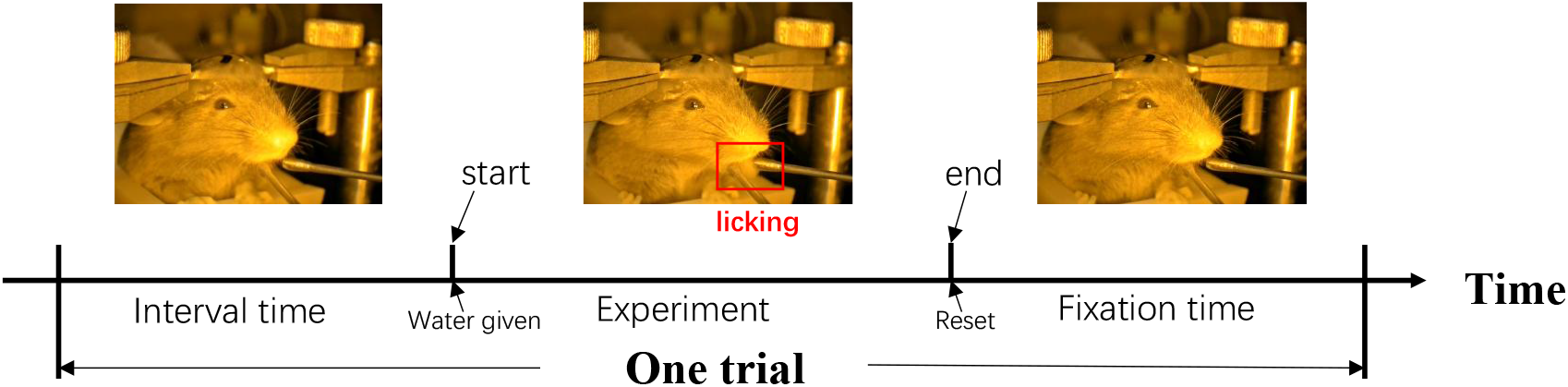
Steps of the experiment procedures.

## Data Records

### Data title

The dataset comprises three brain regions: SNR, VLS, and M2. Within the folder for each brain region, data records are organized by individual mice and recording days. For a specific mouse on a given day, the corresponding folder contains both behavioral data and spike data recorded on that day. Corresponding trials of the data are segmented in behavior file or spike file

### Data content description

All files in our dataset are saved as mat format. To facilitate the investigation of biological mechanisms using this data set, we provide the raw data files along with the detailed experimental protocols described in the articles [17, 18]. These two articles encompass mice behavioral, electrophysiological, and optogenetic stimulation datasets. In this study, we make both raw and preprocessed behavioral data, as well as electrophysiological records, available for public use. Additionally, we include pre-processing and trial segmentation code for the raw data files. This setup allows researchers to freely process the raw data as described in this study and modify them according to their specific research needs.

### Data generation time

The data generation period ranges from December 10, 2019, to December 12, 2020.

### Data volume and data format

The dataset comprises two spike data files: one contains neural activity recorded in the M2 region, and the other includes data from both the VLS and SNR regions. Summary of experimental data statistics is shown in Fig.2. In figure2, we show the summary of data collected during the research procedure. In the figure, x axis is Time and y axis is Mouse ID, different color means different brain regions. We collected data from 20 mice across three brain regions (M2, SNR, and VLS), with a total recording duration of 119 days spanning 203 experimental hours. Each experimental trial comprises 32-channel neural recordings collected throughout the 119-day period, accompanied by corresponding time-series neural spike data.

**Figure 2:**
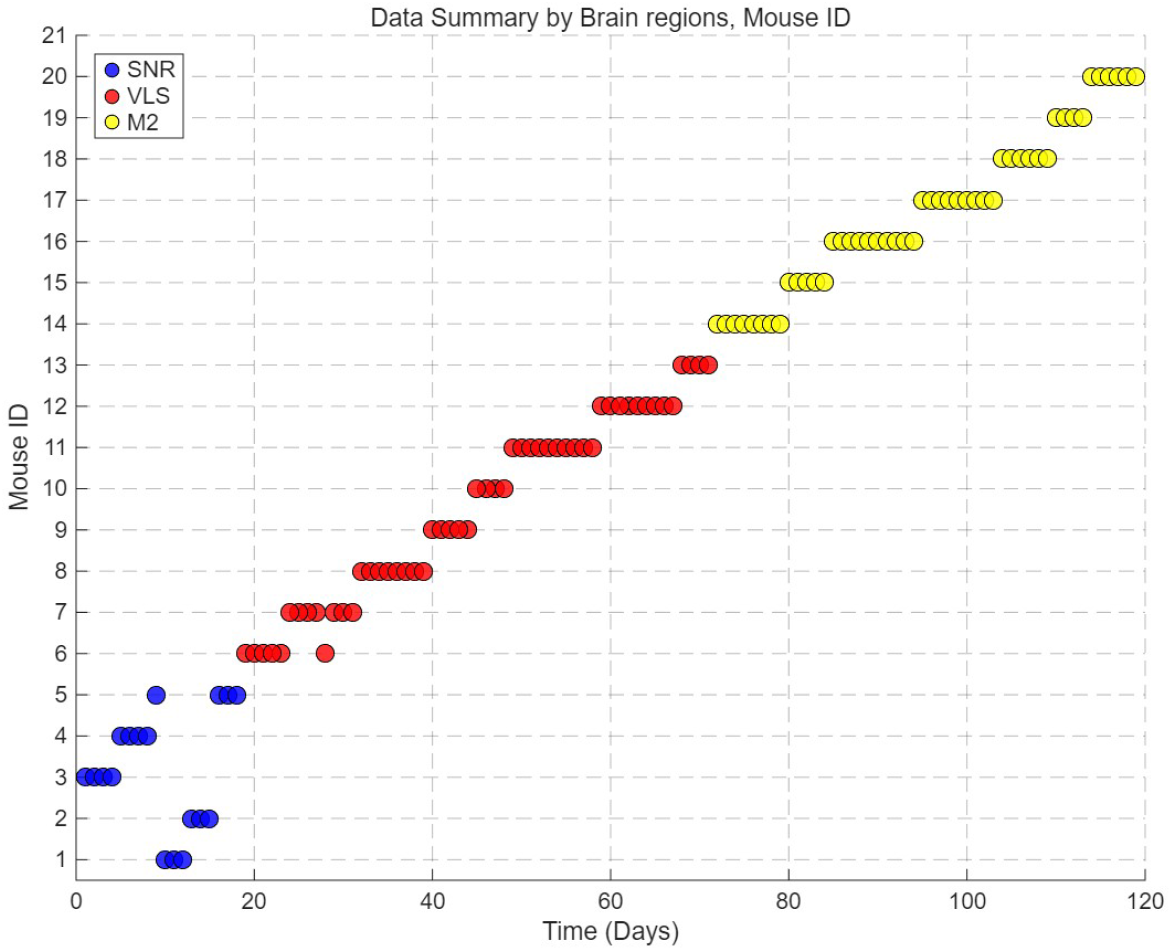
Data statistics summary of the brain regions, mouse ID and experimental days.

Behavioral data are organized in a parallel structure, with separate folders dedicated to the M2 region and the combined SNR/VLS regions. Each folder contains multiple trials documenting lick events in mice, represented as binary values (0 or 1). Behavioral and electrophysiological data were recorded at sampling rates of 2 kHz and 30 kHz, respectively. These datasets can be downsampled to a unified temporal resolution, thereby ensuring precise synchronization between neural spiking activity and behavioral events.

### Data source location

We have published our data set through the open access data repository of the Chinese Academy of Sciences Data Center (https://www.braindatacenter.cn). Moving forward, we plan to release additional data sets tailored for biomedical and computer science research, with the ultimate goal of establishing a comprehensive and systematic data platform.

### Data value

Given the cross-day, cross-subject, and cross-brain-region characteristics of our dataset, researchers can not only employ it for decoding and simulation purposes, but also investigate the biological mechanisms underlying identical behaviors across different brain regions.

### Data generation method

Our research methodology follows a systematic four-stage workflow, as illustrated in Fig.3. The process begins with the simultaneous acquisition of electrophysiological recordings and licking behavior data, followed by precise temporal alignment of neural spike timestamps with licking event markers. The continuous data stream is then segmented into discrete trials corresponding to experimental cycles. The spike data will then be resampled to match the temporal resolution of the behavioral recordings, after which the sampling rate of behavior and spike data will be the same. This comprehensive preprocessing pipeline ultimately yields perfectly synchronized neural and behavioral datasets, ensuring data consistency for subsequent analytical procedures, as shown in Fig.3.

**Figure 3:**
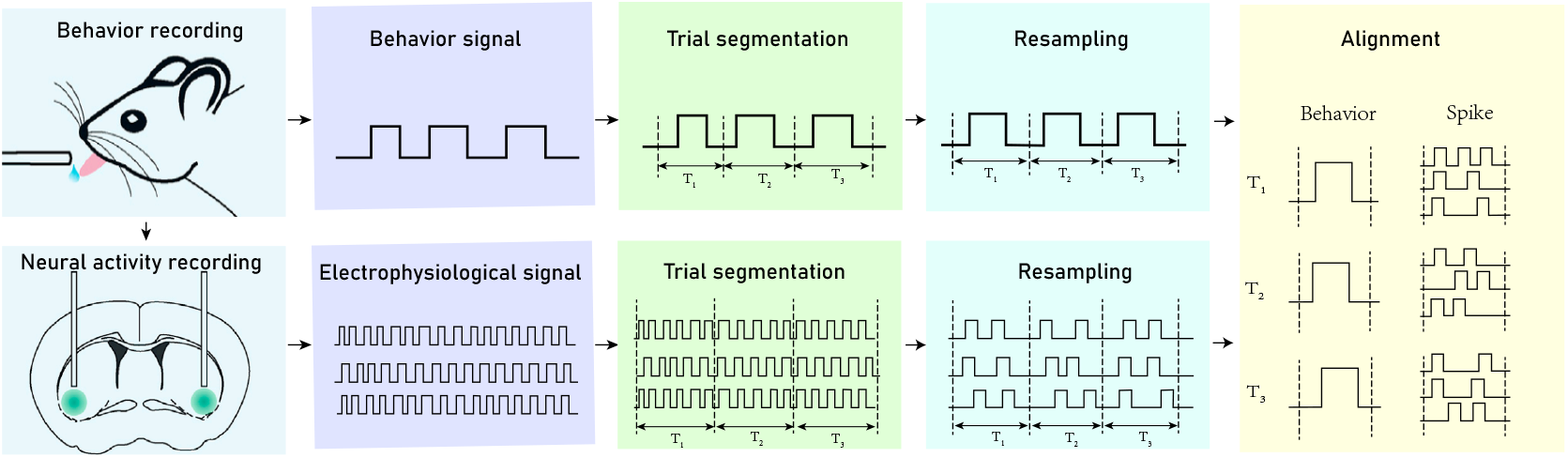
Four steps of data processing: recording, trial segmentation, resampling, and alignment.

Data recording includes two parts: behavioral events and neural signals. Each trial involves synchronized water delivery via a syringe pump, monitoring licking behavior, and recording of neural activity. Behavioral events are digitized as TTL pulses and temporally aligned with neural recordings using the Cerebus acquisition system. Mouse licking behavior is converted to binary time-series data (0 = no lick, 1 = lick) through threshold-based detection to represent event occurrence or non-occurrence respectively. The process is written as Formula 1,

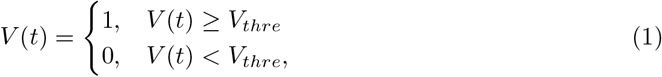

where V(t) represents the amplitude of the behavioral signal at time t and *V*_*thre*_ denotes the detection threshold.

Neural signals are amplified and bandpass-filtered using a Cerebus 32-channel acquisition system (Blackrock Microsystems, UT, USA), with waveforms sampled at 30 kHz. The preprocessing steps for neural spikes consist of the following parts. First, channel selection is performed to exclude channels which failed to collect neuron response by setting their values to zero. Subsequently, spike detection is conducted to identify the timing of neural spikes and each trial is segmented independently. Finally, a resampling procedure is applied to precisely align the sampling rates of neural spiking activity and behavioral data, thereby completing the processing requirements for the AI-ready dataset. Given that different experiments require varying sampling rates, we provide the dataset along with resampling code to facilitate re-sampling for users.

### Data sample description

To demonstrate the correlation between behavior and spikes, we illustrate it, respectively, from M2, SNR, and VLS brain regions in the top left plot of Fig.4. As shown, the horizontal axis represents the trial time (lasting 10 seconds with 20,000 sampling points), while the vertical axis displays the firing of neural spikes and behavior. To facilitate clearer visualization of the results, we applied the resampling function defined in Equation 2 to convert the data into a firing rate time series with a sampling rate of 2 kHz. The consistency between spikes and behavior can be observed in all three regions of the brain, M2, VLS, and SNR. To quantify this relationship, we compute the Pearson correlation coefficient between spikes and behavior. The coefficients exceed 0.7 for all three brain regions, indicating a strong correlation between the data and ensuring its validity for encoding and decoding across regions.

**Figure 4:**
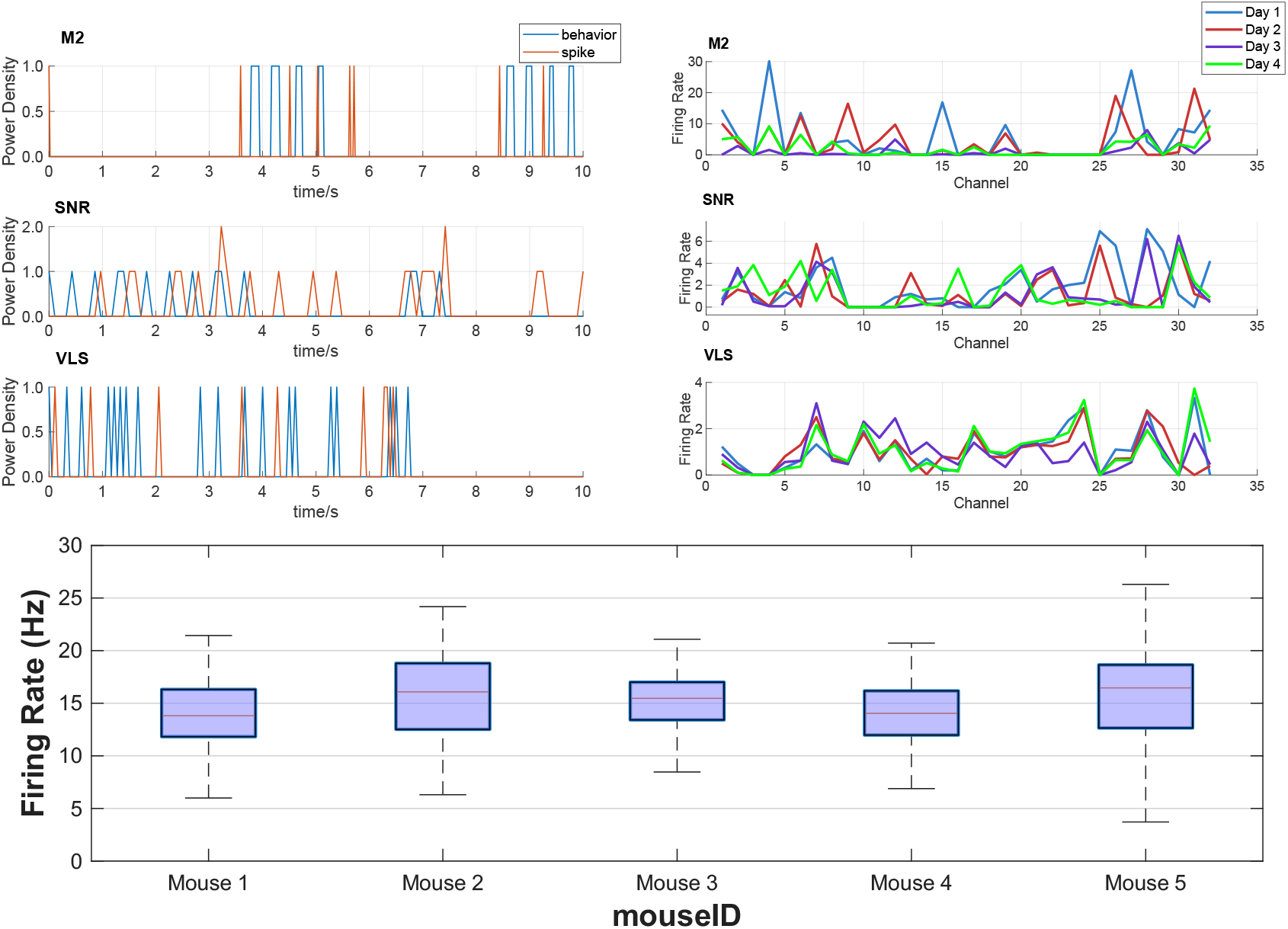
Data consistency of our dataset. The top left plots show consistency of neural spikes and behavior across three brain regions. The top right plots are firing rates across channels through four days. The bottom plots show the box plot of average firing rate distribution across days among different mice in the M2 brain region.

The top right panel of Fig.4 illustrates the variations in firing rates(frequency at which a neuron generates action potentials per second) across different channels of the same mouse over multiple days. The labels M2, SNR, and VLS denote three distinct brain regions. The horizontal axis corresponds to the 32 channels, while the vertical axis indicates the firing rate of each channel. The four polylines represent recordings obtained from the same brain region over four days, with data from M2 collected in one mouse and data from VLS and SNR collected in another. We observed that the distribution of firing rates across channels remains largely consistent over days, exhibiting a Pearson correlation coefficient of 0.76 between different days. This consistency suggests that cross-day decoding for the same mouse is feasible.

The bottom plot in Fig.4 illustrates the distribution of firing rates across different days. The x-axis corresponds to the mouse ID, while the y-axis represents the summed firing rate across all 32 channels. Each box plot displays the average firing rate distribution of one mouse over multiple days. The median firing rates are consistently centered around 15 Hz, with the overall summed firing rates ranging from 10 to 20 Hz. These results indicate the feasibility of cross-subject decoding and simulation.

### Technical validation

To ensure the integrity of the spike dataset, we identify unrelated channels which failed to collect neuron response through offline spike sorting and set them to zero. Signals were sampled at 30 kHz. Spike waveforms were detected by band-pass filtering the data between 250 and 7,500 Hz and applying a threshold set at 3.5 standard deviations (SD) above the background noise level. Subsequent spike sorting was performed offline using the Offline Sorter software. The principal component-based cluster analysis distinguished single units by enforcing: (1) minimum interspike intervals of 1.5 ms and (2) p value for multivariate analysis of variance tests on clusters was *p* ≤ 0.05. Neurons collected from channels that meet any classification criterion are considered lick-modulated, revealing distinct neural populations involved in different aspects of lick sequence control. Following filtering, spike data from relevant cells are recorded. The channels that failed to yield meaningful signals and their corresponding spike data are set to zero. Researchers have shown that excluding cells with a high percentage of coincident spikes can help improve the accuracy of decoding and simulation [19].

To evaluate the practical effectiveness of our dataset, we designed an experiment using 32-channel neural spike data as input to decode mouse licking behavior. We tested the decoding accuracy of this dataset under two algorithmic frameworks: Multilayer Perceptron (MLP) and Support Vector Machine (SVM), with results shown in Fig.5. MLP is a feed-forward neural network capable of learning complex non-linear mappings through multiple hidden layers [20]. SVM is a classical classifier that finds optimal hyperplanes to maximize margin between classes. When using SVM to evaluate the data, we set the preprocessed data as the input for SVM model instead of other features directly. Both are widely used to decode neural signals due to their strong pattern recognition capabilities [21].

**Figure 5:**
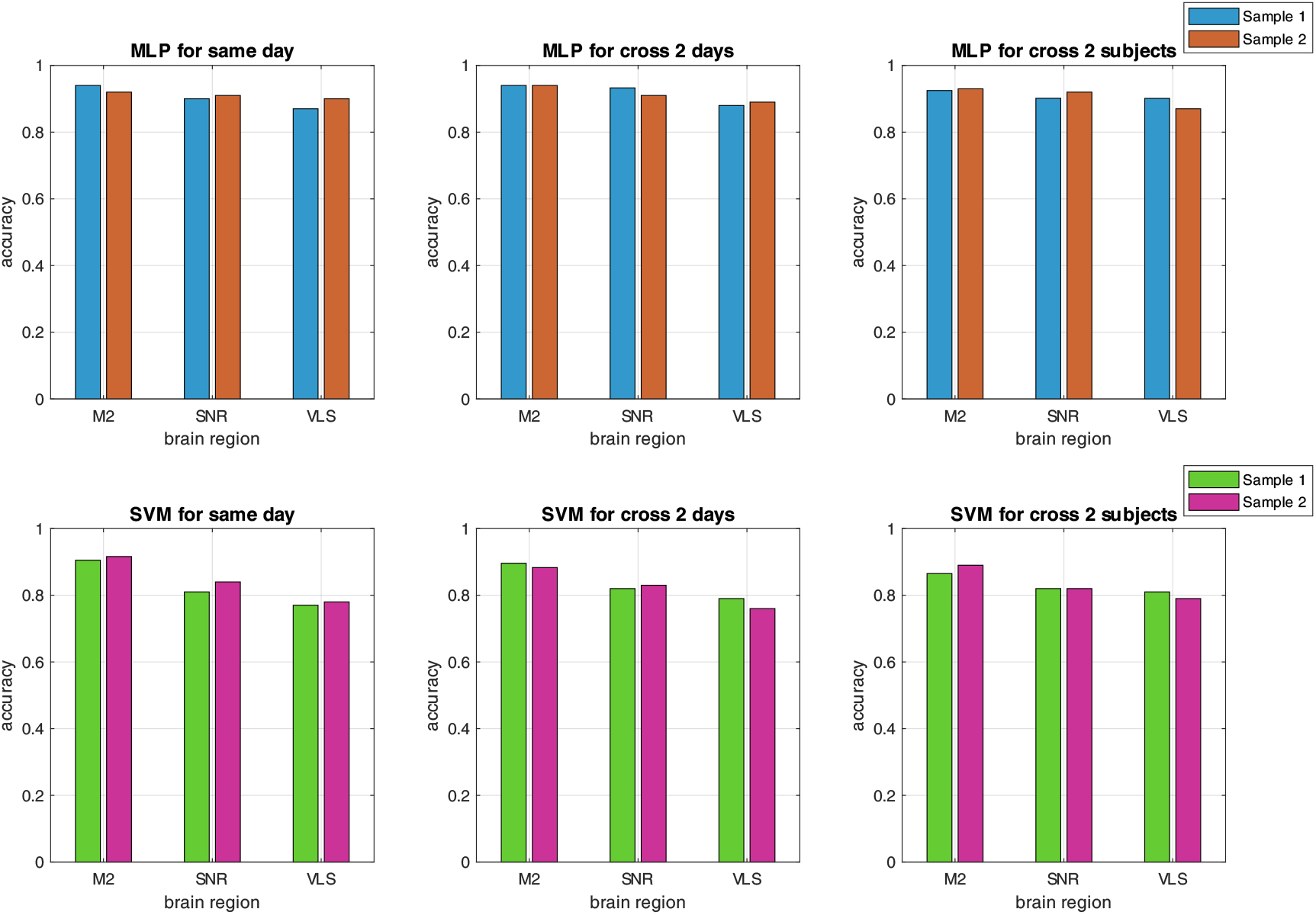
Neural decoding accuracy of MLP and SVM for experimental settings of same-day, cross-2-day and cross-2-subject.

Specifically, Fig.5 illustrates the decoding performance of MLP and SVM algorithms under three experimental conditions. The top panel presents MLP decoding results, and the left graph depicts same-day decoding accuracy across brain regions (represented by dual bars indicating performance from two distinct experimental sessions). The middle graph demonstrates cross-day generalization, where models trained on two daily sessions were evaluated on two other days. The right graph shows cross-subject performance, with models trained on two mice and tested on another two subjects. The bottom panel replicates this experimental paradigm using SVM classification.

Quantitative analysis demonstrates that MLP exhibited superior generalization capabilities, attaining consistently high accuracy in same-day (mean ± SD: 0.94 ± 0.01), cross-day (0.91 ± 0.02), and cross-subject (0.89 ± 0.03) contexts. Additionally, SVM achieved relatively lower decoding accuracy for same-day data (0.90 ± 0.02), with maintained performance under cross-day (0.82 ± 0.01) and cross-subject (0.78 ± 0.02) conditions. These findings indicate that both models effectively utilize the representational capacity of the dataset. The robust performance across both temporal and inter-subject domains confirms the dataset’s utility for neural decoding in scientific research.

### Data usage methods and suggestions

To ensure the alignment between behavior and electrophysiological data, the process of resampling can be implemented for preprocessing:

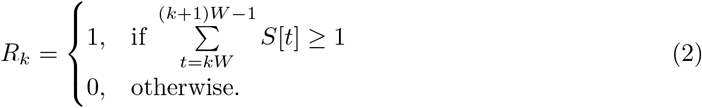

This equation assigns a value of 1 to the resampled output when spikes are detected within the analysis window. The notation uses: *R*_*k*_ (resampling window index), S (sampling points), W (window duration), and t (time). Researchers can modulate the temporal resolution by adjusting the window size parameter W. Some researchers may sum up all spike signals to make the spike and licking water data correspond to each other. The function of this downsampling algorithm can be shown as Formula 3,

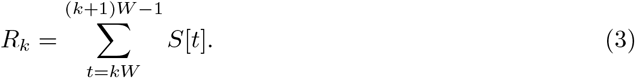

Generally, the resampling process yields synchronized datasets where neural spiking activity and behavioral recordings share identical sampling rates and precise temporal alignment. After resampling the data to the desired sampling rate, this standardized data can be directly applied to investigate cross-day and cross-subject decoding of mouse licking behavior.

The neural spiking data, which is similarly represented as binary events, exhibits a strong temporal correspondence with behavioral states. This alignment enables both behavior decoding (i.e., predicting actions from neural activity) and neural activity simulation (i.e., generating spikes from behavior). The robust correlation between neural firing patterns and behavioral outputs facilitates reliable cross-day and cross-subject investigations using our standardized dataset. For example, the NEDS network [22] was designed to enhance both the encoding and the decoding efficiency simultaneously. Similarly, MiVAE [23] employs a two-level disentanglement strategy to map neural activity and visual stimuli into a shared latent space via artificial neural networks (ANNs), enabling concurrent decoding and simulation within a unified framework.

Spiking neural networks (SNNs), which are event-driven, energy-efficient models using discrete spikes for computation, have been widely adopted across various domains. For instance, SDSA [24] and Semi-SNN [25] have been developed to achieve image classification with reduced energy consumption. SNNs have also demonstrated significant utility in brain-computer interface applications, such as decoding motor activity from electrophysiological signals [26, 27]. Given that our dataset consists entirely of binary events (0s and 1s), it is particularly well-suited for SNN-based research, which not only offers energy efficiency but may also lead to enhanced performance.

## Data Availability

The raw data can be found on the website of the brain data center (https://www.braindatacenter.cn/datacenter/web/#/dataSet/details?id=1722131061926764546 and https://www.braindatacenter.cn/datacenter/web/#/dataSet/details?id=1722129114939236354). The data samples and examples of utilizing samples to implement decoding are specified on a ScienceDB page with DOI (https://doi.org/10.57760/sciencedb.29961).

## Code Availability

The code for importing and processing the data described in this paper, as well as the code for generating the figures and example processing, is available alongside the data in the same public repository (https://doi.org/10.57760/sciencedb.29961).

## Author Contributions

Z.X. co-designed the experiments and developed the code for data processing. B.H. developed the machine learning models discussed in the section of Modeling Performance and co-designed the experiments. L.L. assisted with the data analysis and contributed to the statistical analysis of the dataset. T.X. and Z.C. acquired the data. H.Y. supervised the data collection. T.Z. supervised the overall project, contributed to the study design, and provided critical guidance throughout the experimental and writing phases. All authors contributed to the conception of the study, provided critical feedback on the experimental design and data analysis, and participated in the writing and review of the manuscript.

## Ethics Statement

All animal procedures were conducted according to protocols approved by the Institutional Animal Care and Use Committee (IACUC) of the Institute of Neuroscience, Center for Excellence in Brain Science and Intelligence Technology, Chinese Academy of Sciences (Approval No.NA-013-2019). All the experiment procedure follows the rule of IACUC.

